# Gds1 interacts with NuA4 to promote H4 acetylation at ribosomal protein genes

**DOI:** 10.1101/2021.07.27.454094

**Authors:** Yoo Jin Joo, Stephen Buratowski

**Affiliations:** Department of Biological Chemistry and Molecular Pharmacology, Harvard Medical School, Boston, MA

## Abstract

In our previously published studies, RNA polymerase II transcription initiation complexes were assembled from yeast nuclear extracts onto immobilized transcription templates and analyzed by quantitative mass spectrometry. In addition to the expected basal factors and coactivators, we discovered that the uncharacterized protein Gds1 showed activator-stimulated association with promoter DNA. Gds1 co-precipitated with the histone H4 acetyltransferase NuA4, and its levels often tracked with NuA4 in immobilized template experiments. *GDS1* deletion led to reduction in H4 acetylation in vivo and other phenotypes consistent with partial loss of NuA4 activity. Genome-wide chromatin immunoprecipitation revealed that the reduction in H4 acetylation was strongest at ribosomal protein gene promoters and other genes with high NuA4 occupancy. Therefore, while Gds1 is not a stoichiometric subunit of NuA4, we propose that it interacts with and modulates NuA4 in specific promoter contexts. Gds1 has no obvious metazoan homolog, but structural predictions suggest it may be distantly related to the DEK protein.

## Introduction

Gene expression by RNA polymerase II (RNApII) involves multiple steps and many factors. Transcription is repressed by chromatin occlusion of promoter DNA sequences, so gene activation often requires nucleosome removal or sliding carried out by the Swi/Snf complex and other ATP-dependent chromatin remodelers (1). Subsequent exposure of core promoter sequences allows RNApII and the basal transcription factors to assemble a pre-initiation complex (PIC). This chromatin de-repression is targeted to specific promoters by transcription activators that bind nearby regulatory promoter sequences known as enhancers or Upstream Activating Sequences (UASs). In addition to Swi/Snf and the basal transcription machinery, transcription activators can also recruit the histone acetyltransferases (HATs) SAGA and NuA4 (1, 2). The resulting promoter-localized acetylations are recognized by complexes having one or more bromodomains (3). This group includes Swi/Snf, SAGA, and TFIID, thereby creating a positive feedback mechanism to reinforce acetylation and displacement of the promoter-occluding nucleosomes.

Both histones H3 and H4 are highly acetylated in active promoter regions (4, 5). However, these two histones are modified by distinct HATs: SAGA and NuA3 are responsible for H3 acetylation (H3ac), while NuA4 targets H4 (6). Interestingly, both complexes contain the Tra1 protein, which directly interacts with some transcription activation domains, providing a simple mechanism for coordinated targeting of H3 and H4 acetylation (7). Deletion or point mutations in the H3 and H4 N-terminal tails, where modification sites are concentrated, or their respective HATs, suggest distinct functions for H3ac and H4ac. This difference is presumably because acetylated H3 and H4 tails are “read” by distinct complexes (8). For example, the bromodomain protein Bdf1, a component of both TFIID and the SWR-C complex, preferentially binds acetylated H4 (9, 10), while Swi/Snf recognizes acetylated H3 (11).

We have been studying transcription initiation and elongation using mass spectrometry analysis of transcription complexes assembled on immobilized DNA templates (2, 12–14). In addition to the expected transcription factors, we found that the uncharacterized protein Gds1 was recruited to DNA in an activator-dependent manner. *GDS1* was first identified as a high copy suppressor of slow growth caused by *nam9-1*, a mutant allele of a mitochondrial ribosomal protein gene (15). While Gds1 may therefore have a mitochondrial function, other data suggests a nuclear function. Localization studies show the protein in both the nucleus and cytoplasm (16), and many of the reported physical and genetic interactions of Gds1 are with nuclear proteins ((17–19) https://www.yeastgenome.org/locus/S000005882/interaction). Particularly relevant to this study, Gds1 was identified as a possible substrate for the NuA4 acetyltransferase (20).

Following up on these results, we show here that Gds1 genetically and physically interacts with NuA4. Genome-wide ChIP-seq analysis reveals that Gds1 promotes promoter-proximal H4 acetylation at a subset of genes targeted by NuA4, in particular ribosomal protein (RP) genes and others activated by Rap1. Although there is no obvious metazoan homolog of Gds1, structural predictions suggest Gds1 might be distantly related to metazoan DEK, another poorly understood protein implicated in chromatin regulation of gene expression. While the molecular function of Gds1 remains unclear, our results will help focus future experiments towards solving this mystery.

## Results

### Activator-dependent recruitment of Gds1 to promoters

We have been using an immobilized template assay to isolate and analyze RNApII initiation (2, 12, 13) and elongation complexes (14, 21) in vitro. In these experiments, RNApII PICs were assembled from yeast nuclear extract onto a template containing one Gal4 binding site upstream of the *HIS4* core promoter region (**Fig. 1A**). Digestion with PstI at a unique site upstream of the promoter released downstream DNA template and associated proteins (**Fig. 1B**). As previously reported (2), quantitative mass spectrometry shows Ga14-VP16 activator-stimulated binding of RNApll, the basal initiation factors, Mediator, and the coactivators Swi/Snf, SAGA, and NuA4 (**Fig. 1C**). The fold-enrichment ratios of the coactivators were greater than that of RNApII and its initiation factors, consistent with a basal level of PIC assembly on the naked DNA template.

**Figure 1.**
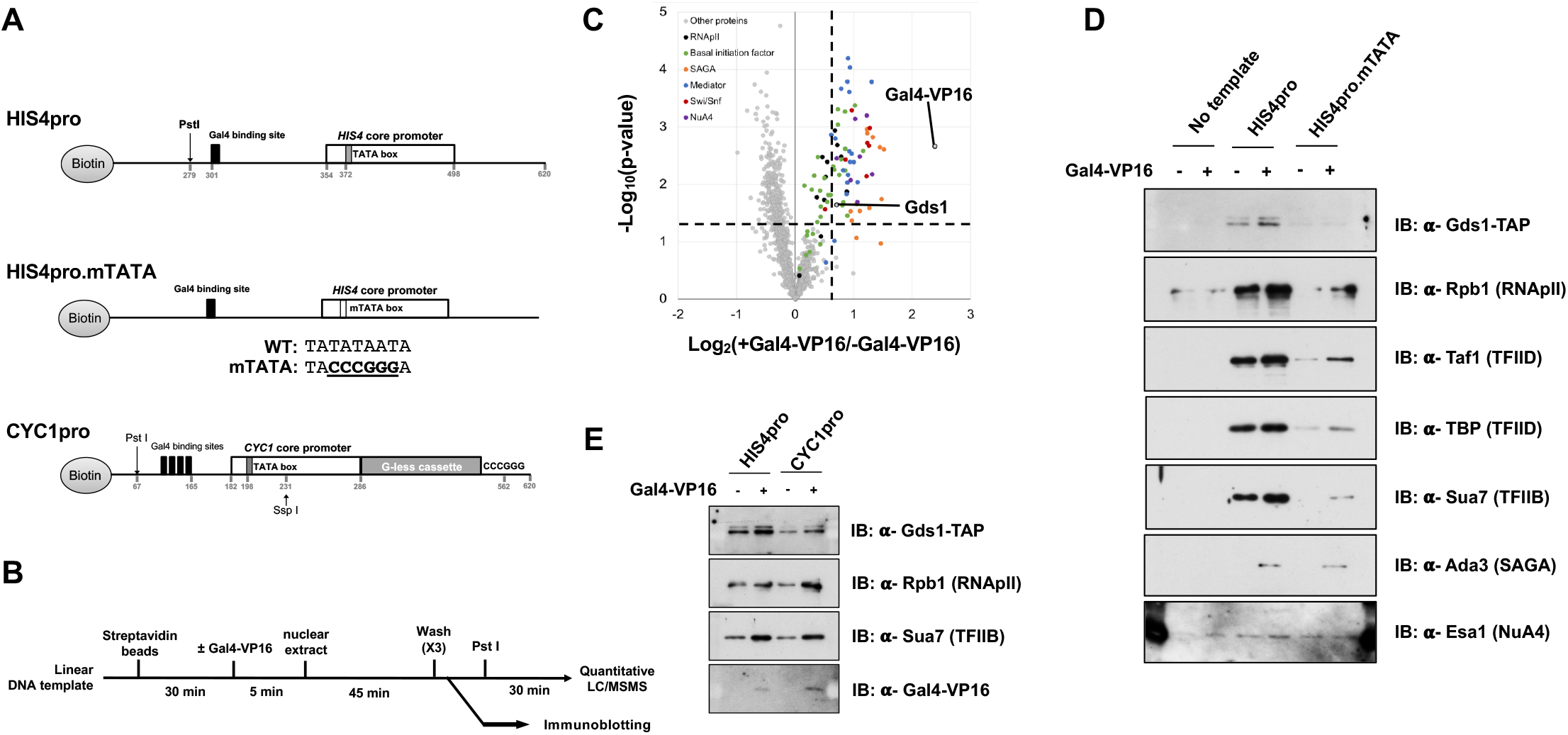
Gds1 is recruited to promoters by the transcription activator Gal4-VP16. (A) Schematic of DNA templates used for immobilized template analysis. Biotinylated fragments were PCR amplified from pSH515 (HIS4pro), pSH514 (HIS4pro.mTATA), or pUC18-G5CYC1 G-(CYC1pro) using oligonucleotides listed in **Sup. Table 2**. (B) Workflow diagram for isolation and characterization of RNApII PICs. (C) Volcano plot showing activator-dependent enrichment (x-axis, vertical dashed line indicates the 95% confidence level using mixed model analysis) of individual proteins on the HIS4pro template versus statistical significance (y-axis, horizontal dashed line shows p-value 0.05). Quantitative mass spectrometry samples from two biological replicates were analyzed with two technical replicates for each. Each dot represents the averaged value for a single protein. Subunits within individual complexes are color coded as indicated. (D) Immunoblots of immobilized template experiments showing that DNA template, transcription activator, and TATA box contribute to PIC and Gds1 binding. Note that SAGA (Ada3) and NuA4 (Esa1) are recruited by activator, but are not affected by the TATA box mutation. (E) Immunoblots of immobilized template experiments showing that Gds1 is recruited to both the *HIS4* and *CYC1* core promoters.

In addition to the expected transcription factors, we also repeatedly observed activator-dependent enrichment of Gds1, a protein of unknown function. To validate the mass spectrometry results, yeast nuclear extract containing TAP-tagged Gds1 was used for PIC formation and immunoblotting assay. Gds1 associated with immobilized transcription template DNA, but not beads alone (**Fig. 1D**). Gds1 binding to DNA was stimulated by Ga14-VP16, and decreased by mutation of the *HIS4* TATA box, similar to the pattern seen for RNApII and several basal transcription factors. NuA4 (Esa1) and SAGA (Ada3) coactivators were also increased by activator, but unaffected by the TATA box mutation. Gds1 association was not specific for the *HIS4* promoter, as similar activator-stimulated binding was also seen on the *CYC1* promoter (**Fig. 1E**). The immobilized templates results suggest that Gds1 may somehow be linked to the RNApII transcription machinery.

### Gds1 interacts with the NuA4 complex

In high-throughput synthetic lethality studies, genetic interactions were seen between *GDS1* deletion and those for factors related to RNApII transcription, RNA processing, DNA replication and repair, and ribosomal biogenesis ((17–19) https://www.yeastgenome.org/locus/S000005882/interaction). Interactors included genes for Eaf1, Eaf3, and Yng2, three non-essential subunits of the NuA4 HAT complex (19). Another study found that Gds1 not only physically interacts with NuA4, but was also a substrate for its lysine acetyltransferase activity (22). We confirmed the interaction between Gds1 and NuA4 by co-precipitation of the NuA4 catalytic subunit Esa1 with Gds1-TAP (**Figure 2A**).

**Figure 2.**
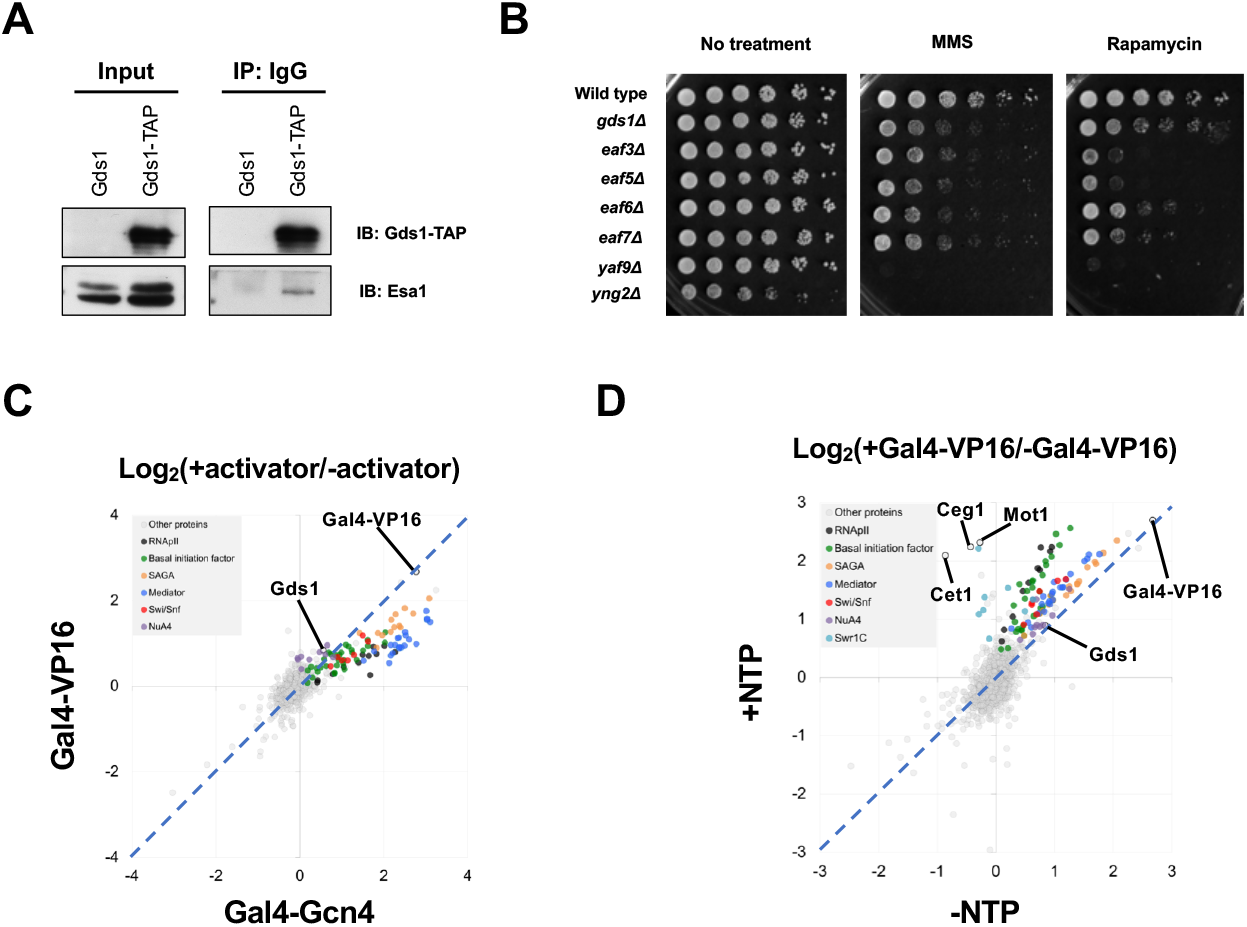
Gds1 interacts with the NuA4 complex. (A) Extracts carrying wild-type or TAP-tagged Gds1 were incubated with IgG beads and pellets were probed for NuA4 subunit Esa1 by immunoblotting. (B) A *gds1Δ* strain was tested alongside NuA4 subunit deletion strains for sensitivity to methyl-methanesulfonate (MMS, 0.02% vol/vol)) or rapamycin (25 mM) by serial dilution spotting. (C and D) Quantitative mass spectrometry shows that Gds1 clusters with NuA4 subunits in differentially associating with VP16 and Gcn4 activation domains (C) and in response to NTPs (D).

To test if Gds1 might have a function related to NuA4, a *gds1Δ* strain was tested for phenotypes seen upon deletion of non-essential NuA4 subunits, including sensitivity to rapamycin and the alkylating agent MMS (23). Like *eaf3Δ*, *eaf5Δ*, *eaf6Δ*, *eaf7Δ*, *yng2Δ*, and *yaf9Δ* strains, a *gds1Δ* strain had slowed growth in the presence of MMS (**Figure 2B**). It also showed sensitivity to rapamycin similar to *eaf5Δ* and *eaf7Δ*, but less so than the other NuA4 subunit deletions.

In our previous immobilized template experiments (2), we found that some coactivators showed differential responses to two different activation domains. SAGA and Swi/Snf were strongly recruited by Gal4 DNA-binding domain fusions to either the VP16 or Gcn4 activation domain. In contrast, NuA4 responded much more strongly to VP16 than Gcn4 (2). In a plot of the mass spectrometry enrichment ratios for these two activators (**Figure 2C**), most activator-recruited factors lie below the diagonal, signifying a stronger response to Gcn4. In contrast, Gds1 clusters with the NuA4 subunits (purple spots) above the diagonal, indicating a preference for the VP16 activation domain. Gds1 also clusters with NuA4 subunits in its response to NTPs (**Figure 2D**). Based on all these observations, we propose that Gds1 interacts with NuA4 in the context of transcription.

### Gds1 affects H4 acetylation at NuA4 target genes

To test for a functional linkage between Gds1 and NuA4 complex, acetylations of histone H3 and H4 acetylation were examined in lysates from wild-type and *gds1Δ* cells. Normalized to total histones (H3 and H2B), H4ac was reduced in the *gds1Δ* cells (**Figure 3A**). In contrast, H3ac was unaffected. This H4ac-specific effect is consistent with a change in NuA4 activity.

**Figure 3.**
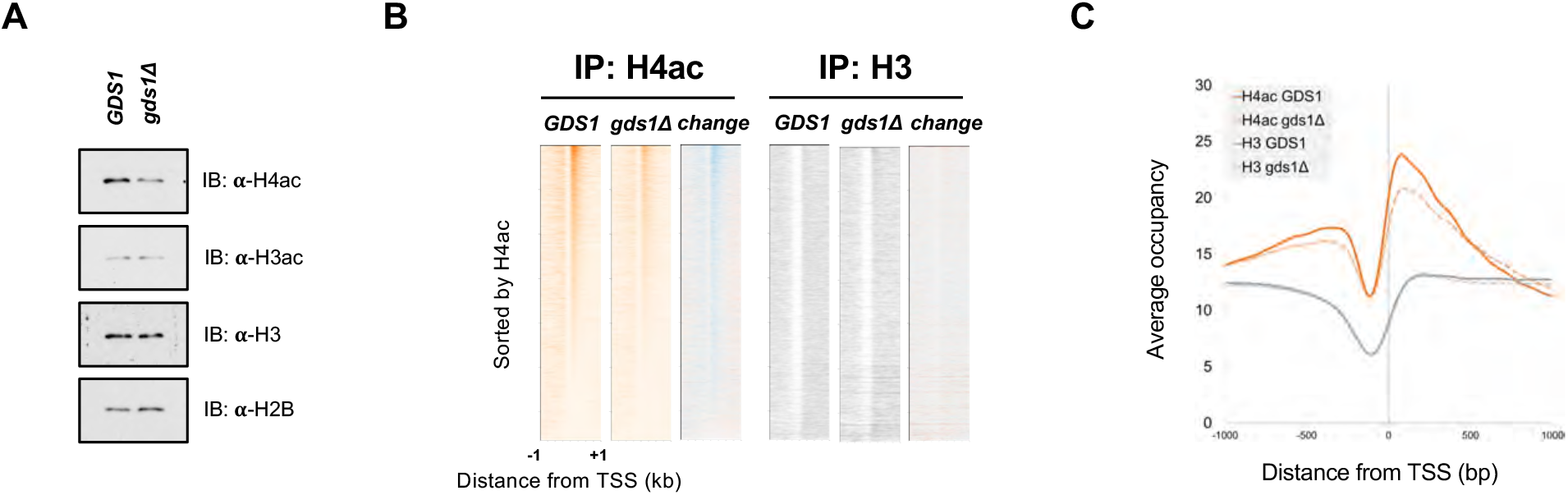
Gds1 promotes histone H4 acetylation. (A) Immunoblotting of whole cell extracts from *GDS1* or *gds1Δ* cells was carried out using the indicated antibodies. (B) Promoter-proximal H4 acetylation was decreased by loss of Gds1. Heat maps show ChIP-Seq analysis for H4ac (orange) and H3 (grey) compared in *GDS1* or *gds1Δ* cells. Each horizontal line represents an individual gene, and genes were ordered by H4ac signal in the *GDS1* strain. For the three panels shown for each immunoprecipitation (IP), the two left panels show the relative levels, while the right panel shows the difference (change = *gds1Δ* – *GDS1*, where blue color intensity represents negative values and orange positive). (C) Metagene analysis of H4 acetylation and H3 data shown in panel B.

To further probe this effect, H4ac was mapped using genome-wide chromatin immunoprecipitation (ChIP-seq). Both heat maps and metagene analysis showed that the H4 acetylation peaks at 5’ regions of most genes were somewhat decreased (**Figure 3B and 3C**). In contrast, total nucleosome occupancy, monitored using anti-H3 antibody, showed no significant change by *GDS1* deletion.

The magnitude of H4ac change for each gene was plotted against its overall acetylation level (**Figure 4A**). Although there was some reduction of H4ac at most genes (median *gds1Δ* /*GDS1* log2 ratio: −0.173), the contribution of Gds1 to acetylation appeared strongest at the most highly acetylated genes. In particular, we found that RPGs (red spots in **Figure 4A**) made up many of the most severely affected genes. Inspection of individual RPG profiles generally showed a pronounced decrease of the H4ac peak overlapping the position of the +1 nucleosome (**Figure 4B**). In many cases, this drop was accompanied by an increase in the total H3 signal at that location, consistent with the role of H4ac in targeting chromatin remodelers and other transcription factors to 5’ ends of genes. The drop in H4ac at RPGs, was verified using standard ChIP-PCR (**Figure 4C and 4D**). Consistent with immunoblotting (**Figure 3A**), H3ac was not similarly affected, with the possible exception of *RPL16A* (**Figure 4D**).

**Figure 4.**
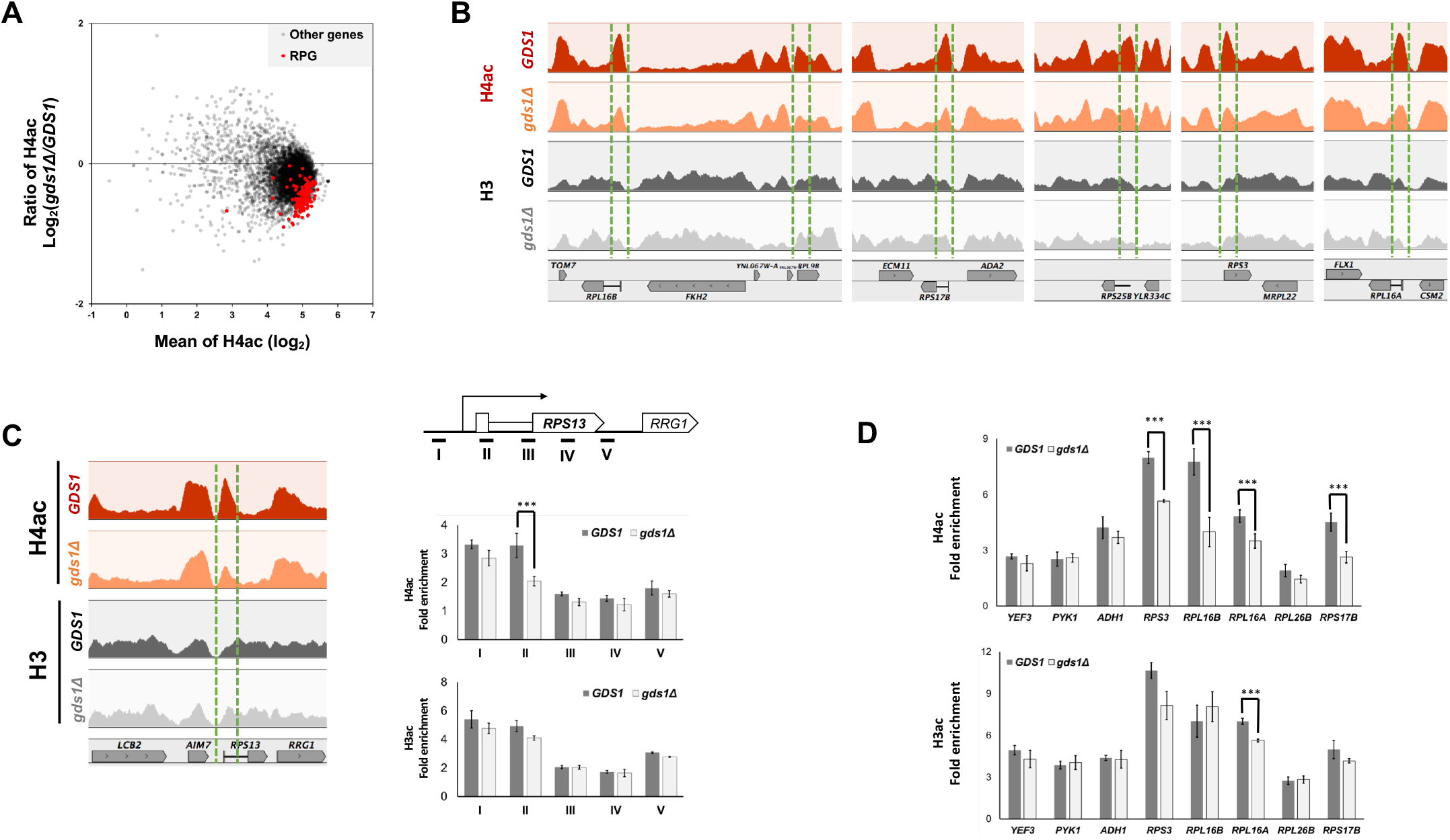
Gds1 stimulation of H4ac is particularly strong at ribosomal protein genes (RPGs). (A) Plot of H4ac levels (log2 of ChIP-seq read counts) versus effect of Gds1 deletion on acetylation (*gds1Δ* /*GDS1* ratio of read counts, expressed as log2). Each spot resembles one gene, and red spots designate RPGs. (B) Browser tracks of H3 (black, *GDS1*; gray, *gds1Δ*) and H4ac (red, *GDS1*; orange, *gds1Δ*) for *RPL9B*, *RPS17B*, *RPS25B*, *RPS3*, and *RPL16A*. Green lines bracket the position of the +1 nucleosomes. (C) Left panel shows browser tracks for *RPS13*, as in panel B. Right panel shows standard ChIP PCR, with schematic of probes at top. Error bars designate standard deviation from three replicates. *** designates P < 0.01. (D) Validation of ChIP-seq data at the 8 indicated promoters, as in panel C.

RPGs are among the most highly expressed and most TFIID-dependent (24–27) genes in yeast. To determine whether either of these two properties correlated with Gds1 function, non-ribosomal protein genes were sorted into quintiles based on their mRNA expression level (**Figure 5A**) or decrease in a *taf1-1* mutant (**Figure 5B**). In both analyses, the change in H4ac levels upon *GDS1* deletion was far less than at RPGs, and there was no statistically significant difference between quintiles. Therefore, Gds1 dependence does not simply correlate with strong expression or requirement for TAF function.

**Figure 5.**
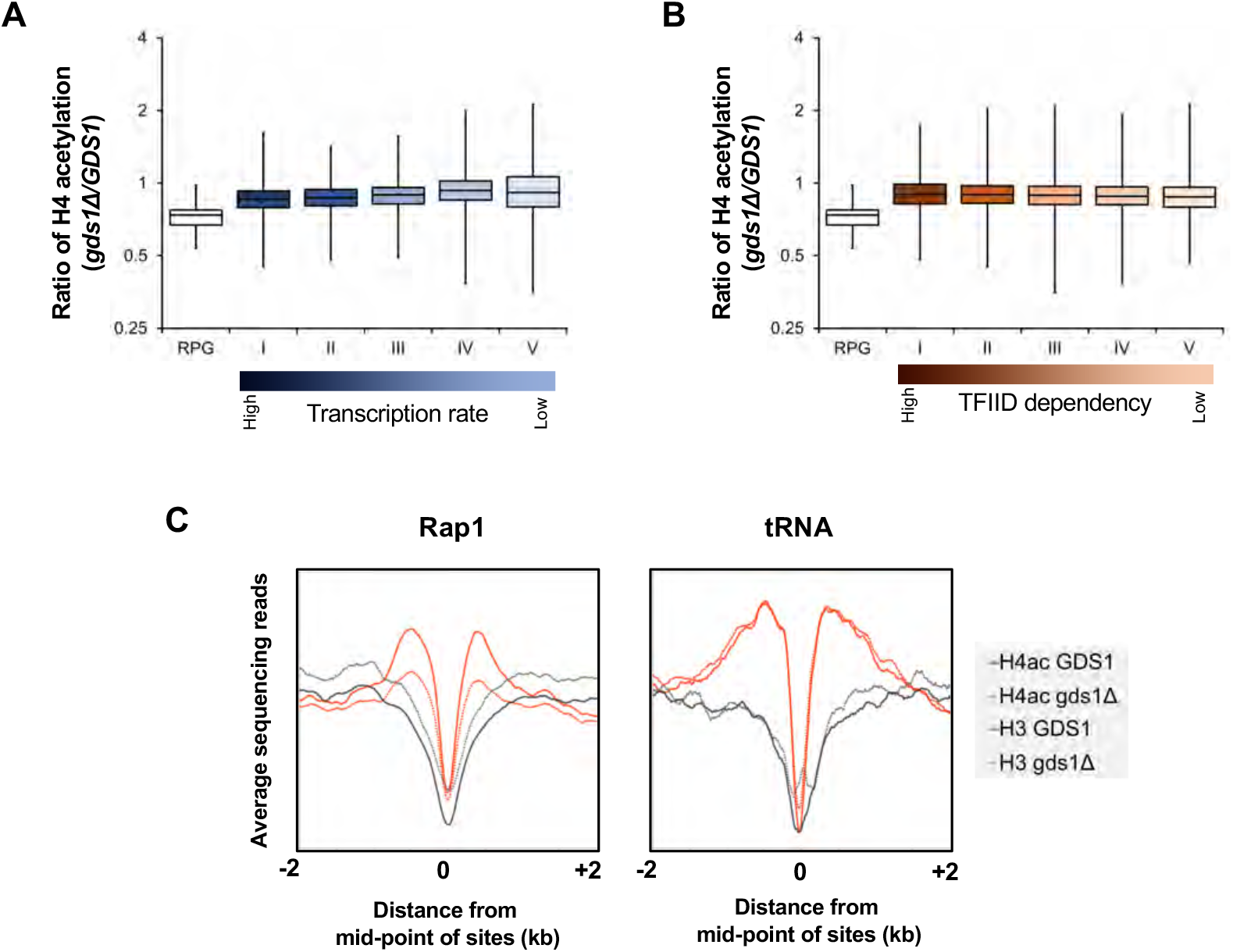
Gds1 effect on H4ac is strong at Rapl binding sites, but is not simply correlated with transcription frequency or TFIID-dependence. (A) Box and whisker plots show the *gds1Δ* /*GDS1* ratio for RPGs, or non-RPGs sorted into quintiles by transcript abundance. (B) Similar analysis, sorting non-RPGs by magnitude of transcription decrease upon TAF1 mutation (24). (C) Metagene analysis for Rapl binding sites (left panel) or tRNAs (right panel), mapping ChIP-seq data from Figure 2.

The transcription factor Rapl binds and activates many RPG promoters. Moreover, Rap1 has been shown to recruit the NuA4 complex, such that high H4ac is observed around Rap1 binding sites (28, 29). A metagene analysis for H3 and H4ac centered around Rap1 binding sites showed a significant decrease of H4ac and increase of H3 in *gds1Δ* cells (**Figure 5C**, left panel). In contrast, no similar effect was seen at tRNA loci, which also have high H4ac (**Figure 5C**, right panel).

We next asked whether response to Gds1 correlated with NuA4 occupancy. ChIP-seq data for the Epl1 subunit of NuA4 (30) was plotted against levels of H4ac (**Figure 6A**). Surprisingly, a strong correlation was not seen. However, RPGs were clearly clustered among genes that had both the highest H4ac and NuA4 occupancy (Figure 6A, left panel, red dots). When the response to *gds1Δ* deletion was mapped on this plot (**Figure 6A**, right panel, color gradient), it was clear that genes with the strongest Gds1 stimulatory effect on H4ac (green and blue dots) were highly enriched among those with highest levels of NuA4. This effect is also apparent in heat maps of individual genes (**Figure 6B**). Plotting the H4ac change for genes with highest levels of Epl1 shows they respond to Gds1 nearly as strongly as RPGs (**Figure 6C**). Based on these results, as well as the experiments above, we propose that Gds1 promotes NuA4 activity at a subset of RNApII-transcribed genes with high NuA4 levels, and particularly at RPGs.

**Figure 6.**
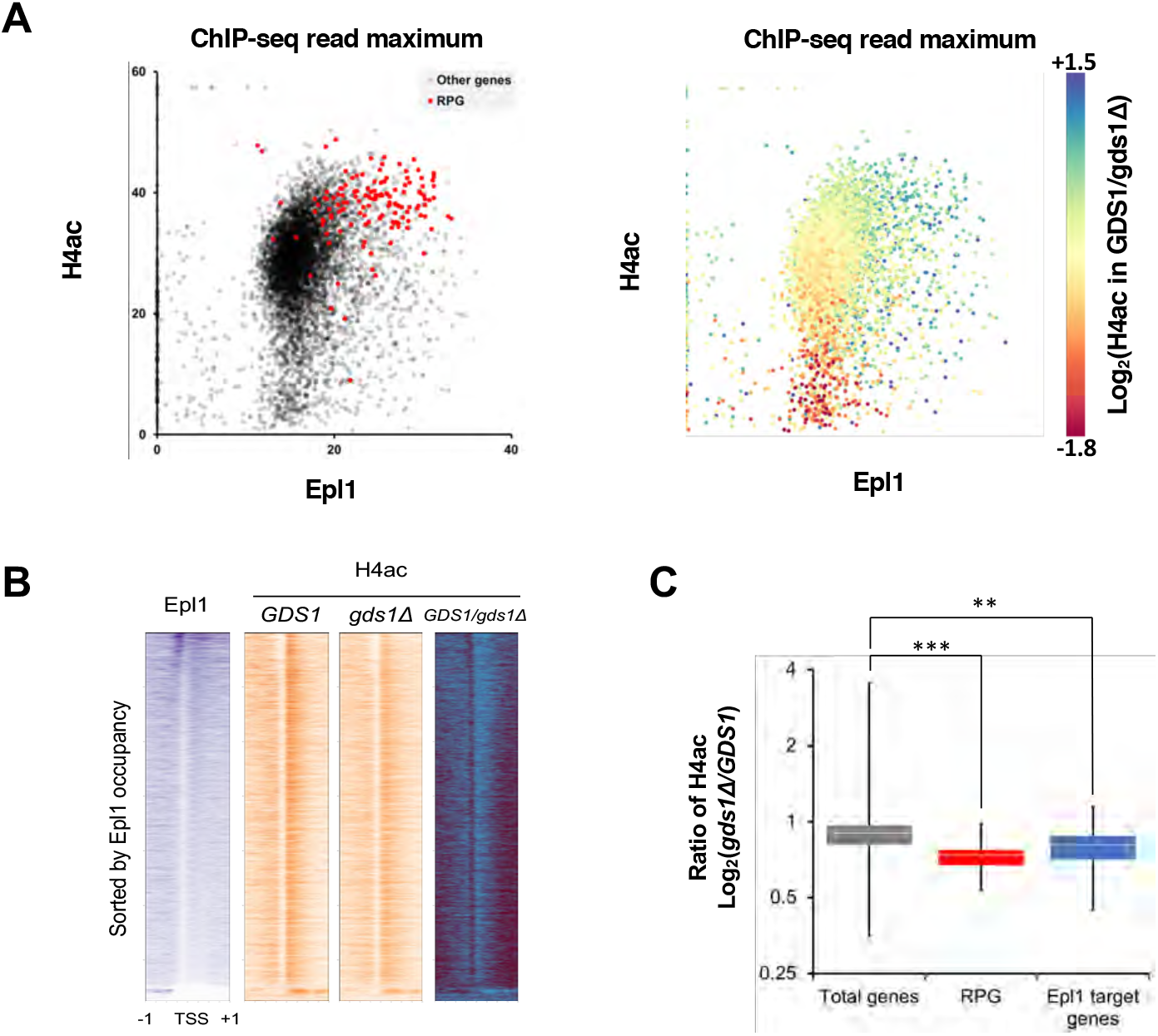
Gds1 effect on H4ac is strongest on promoters with high NuA4 occupancy. (A) Scatter plot of NuA4 occupancy (Epl1 ChIP signal reads (30)) versus H4ac. Left panel shows each promoter as a spot, with RPGs highlighted in red. Right panel shows same data, with effect of Gds1 on H4 acetylation (log2 *GDS1* /*gds1Δ* ratio) color coded. Note that the most Gds1-dependent genes (green-blue) are those with highest Epl1. (B) Heat maps showing H4ac (orange) on individual genes sorted by Epl1 occupancy (purple). (C) Box and whisker plots show the *gds1Δ* /*GDS1* ratio for all genes, RPGs, and Epl1 target genes (defined as the top 4% of genes sorted by Epl1 ChIP signal at the promoter (30)). ** = P<0.05, *** = P< 0.01.

Nuclear extracts were prepared from isogenic *GDS1* and *gds1Δ* strains to test for effects on in vitro transcription. Both extracts were active and responsive to Ga14-VP16 (**Sup Fig 1A**). Linearized HIS4 promoter DNA was assembled into chromatin using recombinant histones and used for immobilized template experiments that included acetyl-CoA (**Sup Fig 1B, C**). No differences were observed in activator-stimulated binding of RNA polII or TFIIH (Kin28). Furthermore, Ga14-VP16 stimulated H4 acetylation, presumably through NuA4 recruitment, but this effect was independent of Gds1 (**Sup Fig 1D**). Therefore, our current in vitro system is unable to reproduce the in vivo effects of Gds1 on histone acetylation, precluding further biochemical analysis.

### Gds1 as a possible yeast ortholog of human DEK protein

Although most NuA4 subunits are conserved over the eukaryotic lineage, a BLASTP search found no metazoan proteins related to Gds1. Homologs were detected among other fungal species, with conservation primarily mapping to a central region encompassing amino acids 90 - 210 (**Sup Fig 2A**). Given that conserved regions often correspond to a structural unit, secondary structure prediction and 3D modeling of the conserved central domain was performed using Phyre2 (31), Robetta (32), and I-TASSER (33). All three programs predicted a domain of four or five alpha helices, with the three longest forming a 3-helical bundle (**Sup Fig 2B, C**). Interestingly, this predicted domain scored as a strong structural match for the C-terminal domain of the mammalian DEK protein (**Figure 7A and 7B**), which has been implicated in chromatin structure and gene expression (34, 35). The DEK-C fold is found in the DNA binding domains of E2F/DP transcription factors, but also in chitin synthetases and some kinases, so its exact function remains unclear.

**Figure 7.**
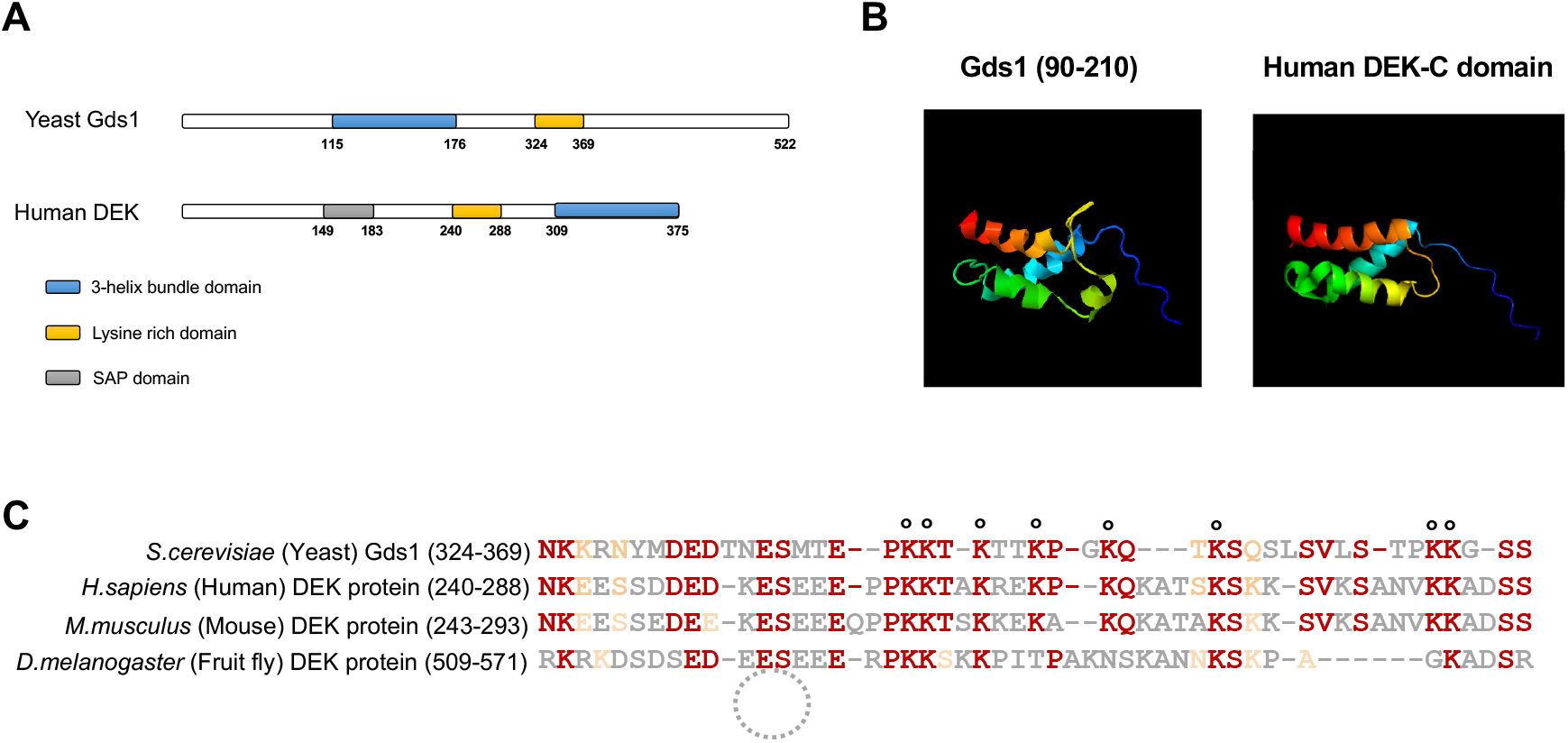
Gds1 is a potential yeast ortholog of metazoan DEK protein. (A) Schematics of predicted domain structures of Gds1 and human DEK protein. (B) Predicted structure for the Gds1 region most highly conserved in other yeast homologs (left panel, see **Supp. Figure 1**), compared to the DEK-C structure: PDB entry 1Q1V (43). (C) Sequence alignment of the Gds1 lysine-rich region with a lysine-rich region from human, mouse, and fruit fly DEK. Circles mark NuA4 acetylation target sites identified in Gds1 (20).

Regions of Gds1 outside the helical domain are enriched for charged and polar residues, and are predicted to be disordered. DEK also has many regions of low complexity, and pair-wise comparison of the two proteins shows both contain a lysine-rich domain that can be aligned, albeit with low statistical significance (**Figure 7C**). Interestingly, lysine residues within this region of Gds1 are acetylated by NuA4 in vitro, although the functional relevance of these modifications is unclear (20). In our immobilized template experiments, the absence or presence of acetyl-CoA did not change the amounts of Gds1 bound to promoters, arguing that acetylation is not required for Gds1 binding (**Sup Fig 1E**).

As DEK is believed to function in transcription regulation through its interactions with chromatin (34, 35), and given the possible structural similarity, it is tempting to speculate that Gds1 could be a possible yeast ortholog. When DEK was expressed in yeast lacking Gds1, it was unable to complement the MMS sensitivity of *gds1Δ* (data not shown). However, yeast-human cross-species complementation by known orthologs is generally possible for only the most highly conserved proteins, so a definitive answer will have to wait until the functions of these two proteins are better understood.

## DISCUSSION

Genetic and biochemical studies have revealed a surprisingly large number of basal factors, co-activators, and chromatin-related proteins needed for activator-stimulated transcription. To study these factors as integrated system, we have been using yeast nuclear extracts to assemble complexes on both naked DNA and chromatin templates (2, 12–14). Our proteomic analyses of initiation and elongation complexes suggest nuclear extracts effectively reproduce many aspects of in vivo transcription. In addition to all the expected co-activators and basal transcription factors, transcription on immobilized templates also recapitulates the cycle of RNApII phosphorylations driving the exchange of elongation and mRNA processing factors (14). Few unexpected or unidentified proteins were similarly enriched, suggesting that most or all of the key factors have been discovered.

One exception is Gds1, a poorly characterized protein that reproducibly showed activator-stimulated association with transcription templates (**Fig. 1**). *GDS1* was first identified as a high copy suppressor of *nam9-1*, a mutation in a mitochondrial ribosomal protein that cannot grow on media containing glycerol as a carbon source (15). Although Gds1 protein can be detected in mitochondria (36), it is predominantly in the cytoplasm and nucleus (16). A *GDS1* deletion shows synthetic sick and lethal phenotypes with many nuclear proteins linked to transcription and ribosome biogenesis, including components of the NuA4 HAT complex ((17–19) https://www.yeastgenome.org/locus/S000005882/interaction).

The results presented here further connect Gds1 with NuA4. We confirm (**Fig. 2A**) an earlier report that Gds1 co-precipitates with NuA4 (22), and show that loss of Gds1 reduces H4 acetylation in vivo (**Figs. 3 and 4**). Although Gds1 is not an integral subunit of NuA4 (37), we find that it often clusters with NuA4 subunits during quantitative mass spectrometry of transcription complexes (**Fig. 2B, C**). The drop in H4ac upon *GDS1* deletion correlates with high NuA4 occupancy, but not all promoters with high H4ac levels depend on Gds1 (**Figs 3B and 4A**). Therefore, Gds1 may regulate NuA4 activity in a manner sensitive to promoter context. Consistent with this possibility, Gds1 binding to immobilized templates was increased by the presence of the TATA-containing core promoter, in this respect resembling PIC components more than NuA4. Unfortunately, while our in vitro transcription system exhibits activator-stimulated H4 acetylation, it apparently does not reproduce the stimulatory effect of Gds1 (**Sup Fig 1**).

We found that ribosomal protein gene promoters, which typically have some of the highest histone acetylation levels, are highly enriched among the promoters most strongly affected by loss of Gds1. RPGs comprise a unique regulon designed to tune mRNA and protein expression to the available amounts of the RNA polI-transcribed rRNA (38). This balance is sensitive to availability of nutrients and cellular growth and division rates. This signaling may even underlie the original genetic interaction between *gds1Δ* and a mitochondrial ribosomal protein mutation. However, the Gds1 effect on RPG H4ac is not simply an indirect effect of slower growth, as *gds1Δ* cells grow at normal rates on rich media (**Fig. 2B**).

Future experiments will undoubtedly reveal more about how Gds1 fits into gene regulation. Searches for homologs that might supply further clues were mostly uninformative. However, structural prediction programs produced a limited, but intriguing, potential similarity between yeast Gds1 and metazoan DEK protein. DEK is also a small protein implicated in gene expression and other nuclear functions. Like Gds1, its mechanism of function remains mysterious, but it is notable that DEK associates with genomic regions enriched for H4 acetylation (39). Furthermore, it has been reported that

DEK can influence the activity of the p300 and PCAF HAT complexes (40). While a link between Gds1 and DEK is purely speculative at this point, it is worth consideration given the dearth of other clues to their functions.

## Acknowledgements

We are grateful to Craig Mizzen, Jacques Côté, Tony Weil, and Joe Reese for antibodies, and Steve Hahn for transcription template plasmids. This work was supported by NIH grant GM046498 to S.B.

## MATERIALS AND METHODS

### Strains, plasmids, and oligonucleotides

*S. cerevisiae* strains, oligonucleotides, and plasmids used in this study are listed in **Supplemental Tables S1, S2, S3.**

### Yeast techniques

For phenotypic analyses, overnight liquid cell cultures of the indicated yeast strains were normalized for cell density, serially diluted (3-fold in each step), and spotted onto the indicated media.

Whole cell extracts were made by glass bead lysis as previously described (41). For testing protein interactions, IgG-agarose beads were added to bind Gds1-TAP, and after washing, associated proteins were analyzed by elution in loading buffer. For all immunoblotting, proteins were separated by SDS-10% polyacrylamide gel electrophoresis, transferred to Immobilon P membrane (Millipore Sigma), immunoblotted with the designated primary antibodies (listed in **Supplemental Table 4**), and the appropriate secondary antibody conjugated with horseradish peroxidase. Detection was by chemiluminescence using Pierce Supersignal West detection reagent.

### Immobilized template binding assay

Immobilized templates experiments were carried out using yeast nuclear extracts as previously described (2, 13, 14, 21). Briefly, the indicated DNA templates were prepared by polymerase chain reaction, with one primer containing a biotin residue for linkage to magnetic beads (Invitrogen MyOne Strep Dynabeads). Immobilized templates were incubated with yeast nuclear extracts, followed by washes to remove unbound proteins, and isolation of template-bound proteins using magnetic concentration. To analyze bound proteins by standard immunoblotting, the beads were resuspended in loading buffer and loaded on to SDS-polyacrylamide gels. Antibodies used are listed in **Supplemental Table 4**. For quantitative mass spectrometry, proteins were eluted from beads using Pst I and subjected to multiplexed iTRAQ or TMT labeling, followed by 3D-LC/MS-MS analysis. Note that the quantitative mass spectrometry figures in this paper show re-analysis of data originally generated in Sikorski et al. (**Fig 2B**, (2)) and papers by Joo et al. (**Fig 2C**, (14, 21)).

### Chromatin immunoprecipitation and ChIP-Seq

Chromatin immunoprecipitations were performed using antibodies against histone H3 (Abcam ab1791) and acetylated histone H4 (Millipore 06-598). Genome-wide data was generated and analyzed as previously described (13, 42), and has been deposited in GEO under accession number XXXX. For gene-specific ChIPs, standard PCR was performed using primers listed in **Supplemental Table S2**.

## Supplemental Information for Joo and Buratowski

**Supplementary Figure 1.**
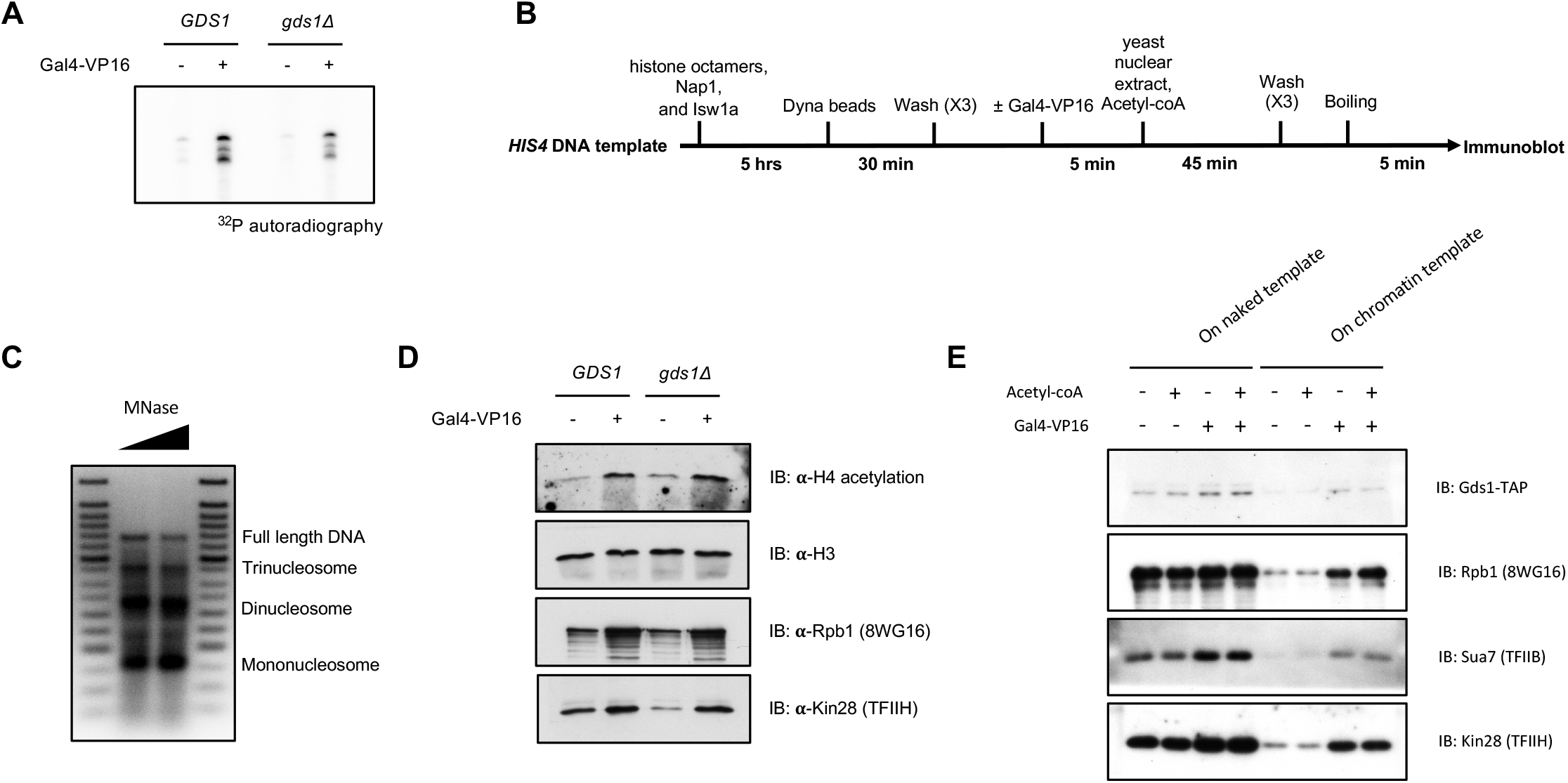
In vitro transcription and histone acetylation using *gds1Δ* nuclear extract. (A) In vitro transcription reactions performed with *GDS1* or *gds1Δ* nuclear extracts, in the presence (+) or absence (-) of Ga14-VP16 structure. (B) Workflow diagram for isolation and characterization of RNApll PICs on chromatinized templates. (C) Micrococcal nuclease (MNase) analysis of chromatinized templates showing protected DNA bands corresponding to the indicated nucleosome species. (D) Immunoblotting of proteins associated with immobilized chromatin templates analyzed as shown in part B, comparing *GDS1* or *gds1Δ* nuclear extracts. Note the increase H4 acetylation in the presence of activator. (E) Immunoblotting to test how chromatin and acetylation (+Acetyl-CoA) affect activator-stimulated Gds1 recruitment.

**Supplementary Figure 2.**
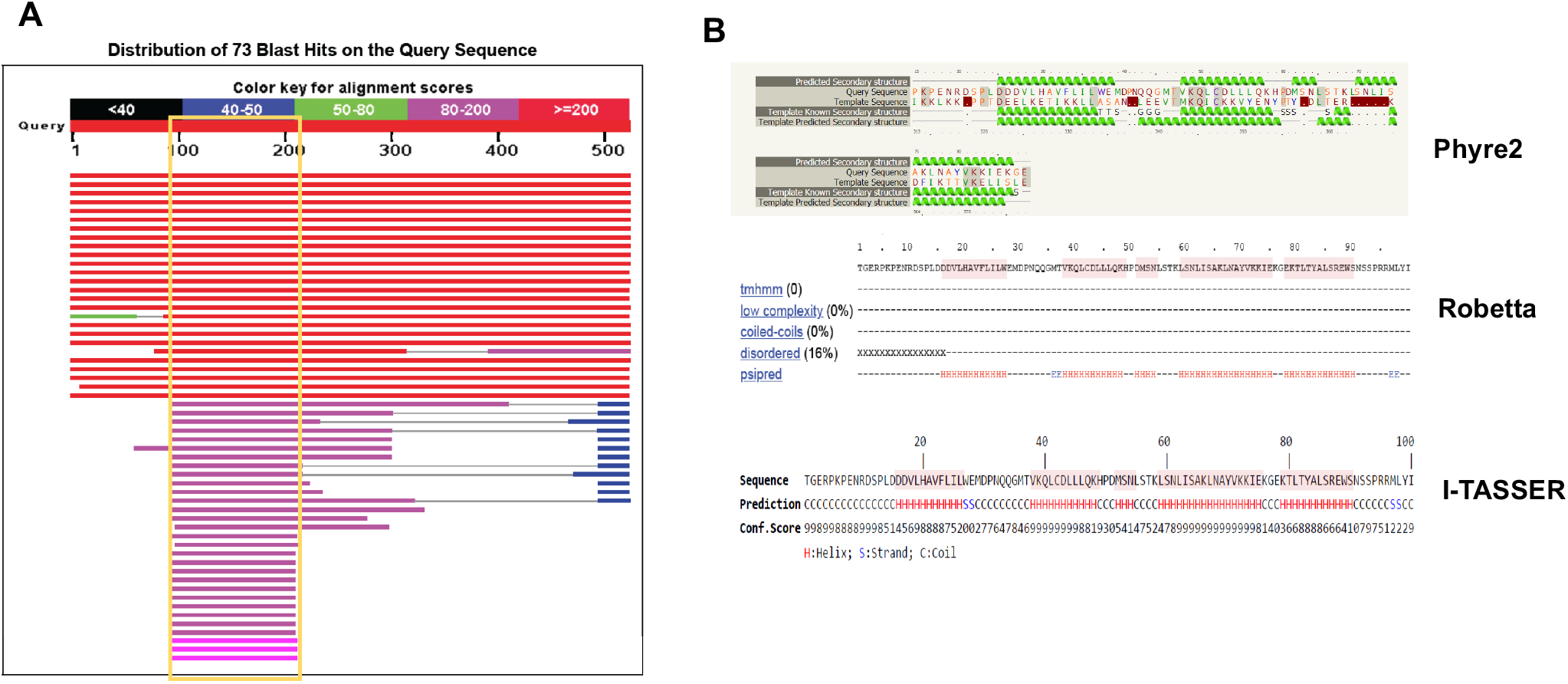
Prediction of Gds1 domain structure. (A) Schematic of BLAST results showing that amino acids 90-210 comprise the most highly conserved region in Gds1 homologs. (B) Protein structure predictions of Gds1 90-210 using Phyre2 (31), Robetta (32), and I-TASSER (33).

**Supplemental Table 1:**
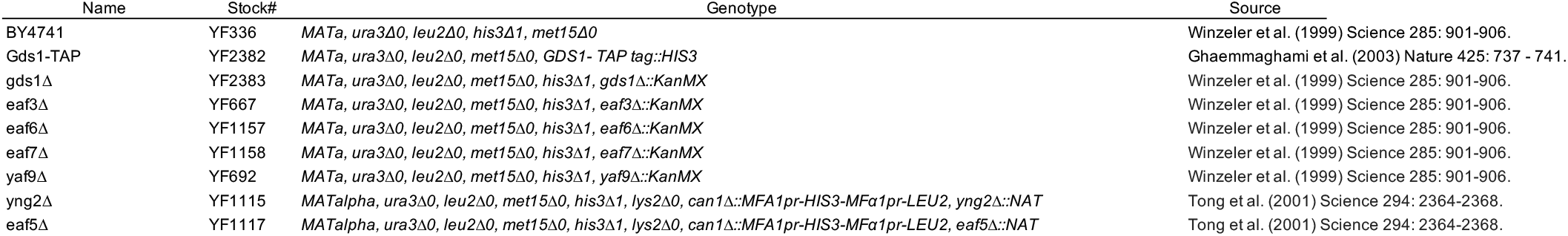
Yeast strains used in this study.

**Supplemental Table 2:**
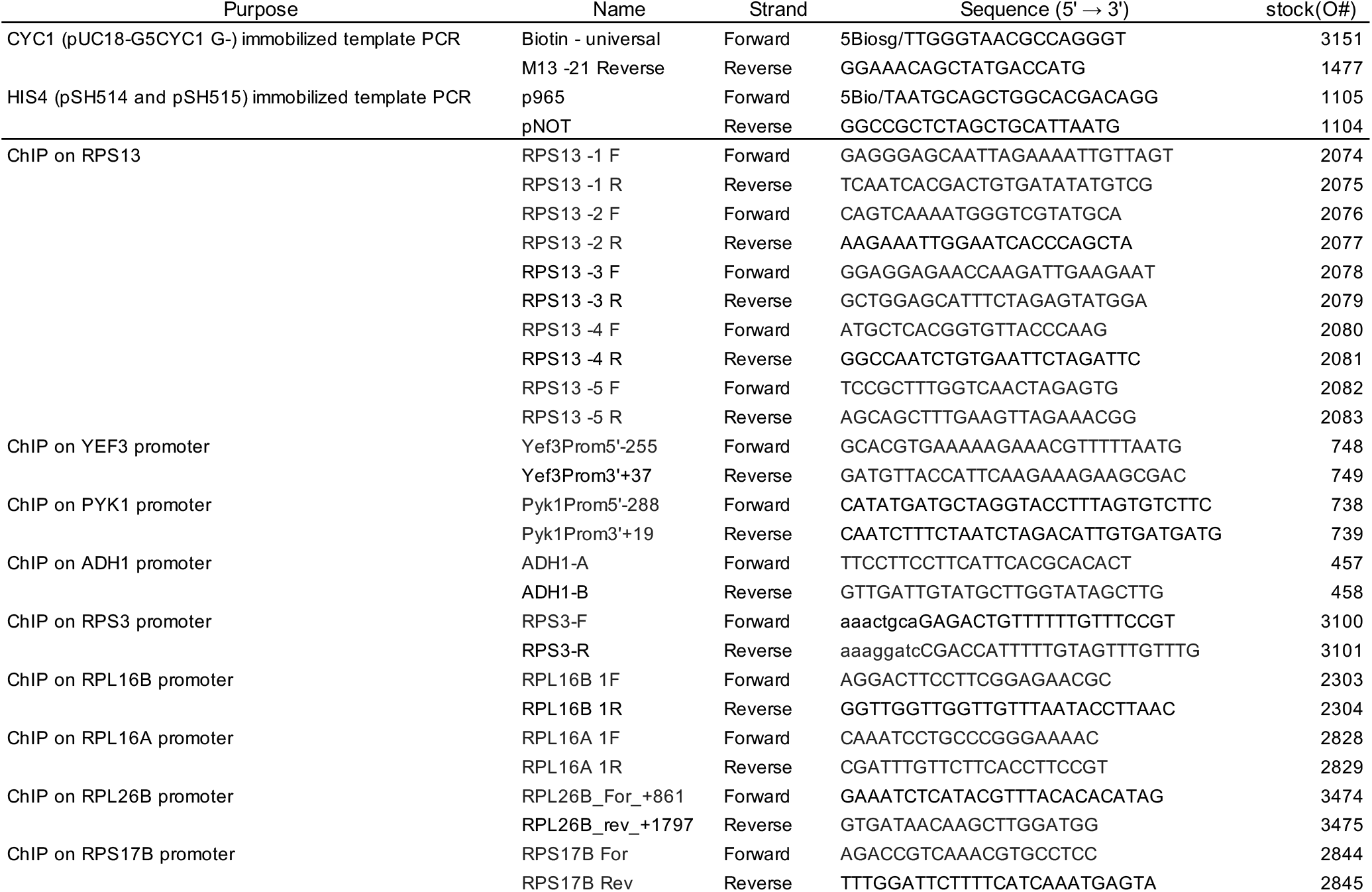
Oligonucletides used in this study.

**Supplemental Table 3:** Plasmids used in this study.

| Plasmid name | Stock # | Features | Source |
| --- | --- | --- | --- |
| pUC18-G<sub>5</sub>CYC1 | SB649 | 5 copies of GAL4 binding sites upstream of *CYC1* promoter driving short G-less cassette (250, 277 nt tra | Johnson et al. (2009) Mol. Cell 35: 769-781. |
| pSH515 | F529 | HIS4 core promoter region with a single Gal4 binding site, AmpR | Ranish et al. (1999) Genes & Dev. 13: 49-63. |
| pSH514 | F529 | HIS4 core promoter region with a mutated TATA box and single Gal4 binding site, AmpR | Ranish et al. (1999) Genes & Dev. 13: 49-63. |

**Supplemental Table 4:** Antibodies used in this study.

| Protein recognized | Complex | Antibody type | Source |
| --- | --- | --- | --- |
| Histone H3 | Nucleosome | Rabbit polyclonal | Abcam ab1791 |
| Histone H2B | Nucleosome | Rabbit polyclonal | Abcam ab1790 |
| Histone H4ac | Nucleosome | Rabbit polyclonal | Millipore 06-598 |
| Histone H3ac | Nucleosome | Rabbit polyclonal | Millipore 06-599 |
| TATA Binding Protein | TFIID | Rabbit polyclonal | Buratowski lab |
| Kin28 | TFIIH | Rabbit polyclonal | Babco PRB-250C |
| Sua7 | TFIIB | Rabbit polyclonal | Buratowski lab |
| Rpb1 | RNA pol II | Mouse monoclonal 8WG16 | Buratowski lab |
| Esa1 | NuA4 | Rabbit polyclonal | Craig Mizzen |
| Taf1 | TFIID | Rabbit polyclonal | Tony Weil |
| Ada3 | SAGA | Rabbit polyclonal | Tony Weil |
| TAP tag | Used for Gds1-TAP | Rabbit polyclonal | Peroxidase anti-Peroxidase antibody Sigma P1291 |
| His6 | Used for Gal4-VP16 | Mouse IgG1 monoclonal 4A12E4 | ThermoFisher 6x-HIS Tag monoclonal antibody 37-2900 |

